# Bounded environmental stochasticity generates secondary Allee thresholds

**DOI:** 10.1101/2025.04.03.647087

**Authors:** Sebastian J. Schreiber

## Abstract

A population exhibits an Allee effect when there is a critical density below which it goes extinct and above which it persists. Classical models with environmental stochasticity predict inevitable extinction, stemming from the assumption that environmental variation is normally distributed with rare but arbitrary large effect sizes. However, environmental fluctuations are bounded and often not normally distributed. To address this reality, I analyze piecewise deterministic Markov models (PDMPs) of populations experiencing Allee effects, where environmental dynamics are governed by a finite-state Markov chain. These models predict that populations can persist through the emergence of two threshold densities. Below the lower threshold, populations deterministically go extinct; above the higher threshold, they deterministically persist. At intermediate densities, populations experience stochastic bistability: with positive, complementary probabilities, they either go extinct or persist. Persistence becomes impossible when the carrying capacity in one environment falls below the Allee threshold in another. Such mismatch occurs only when the environmental state affects percapita growth rates non-monotonically, as when environments supporting higher carrying capacities also produce higher predation levels or greater mate limitation. This work demonstrates that incorporating realistic bounded environmental fluctuations substantially alters predictions about population persistence, with important implications for conservation and management.

## Introduction

Populations exhibit an Allee effect when, at low densities, the per-capita growth rate increases with density (Allee, 1931; Stephens et al., 1999). Common causes of this effect include larger population sizes that reduce the risk of individual predation and the difficulty of finding mates, or increase cooperative foraging or breeding (Courchamp et al., 1999; Stephens et al., 1999; Gascoigne and Lipcius, 2004; Courchamp et al., 2008; Gascoigne et al., 2009; Kramer et al., 2009). When an Allee effect is sufficiently strong, it results in a critical density below which a population is driven rapidly to extinction and above which the population grows and persists. This critical density can play an important role in the rescue of threatened populations (Dennis, 1989; Stephens and Sutherland, 1999; Berec et al., 2007; Courchamp et al., 2008), controlling diseases or pest species (Hilker et al., 2009; Tobin et al., 2011; Kang and Castillo-Chávez, 2014), or managing harvested populations (Berec et al., 2007; Winter et al., 2019).

Populations that simultaneously experience a strong Allee effect and environmental stochasticity may experience a greater risk of extinction (Dennis, 1989, 2002; Lande et al., 2003). Stochastic fluctuations in environmental conditions can knock a persisting population below the Allee threshold and initiate an exponential decline to extinction. Traditional models of environmental stochasticity have predominantly used diffusion processes or stochastic differential equations (SDEs) (Dennis, 1989, 2002; Lande et al., 2003). These approaches assume that environmental fluctuations are normally distributed where small deviations from the mean occur with the greatest likelihood but where arbitrarily large deviations are possible (Dennis, 1989, 2002; Liebhold and Bascompte, 2003; Lande et al., 2003; Dennis et al., 2015).

An important consequence of this unbounded stochasticity is that populations with a strong Allee effect have no stable non-zero stationary distribution (Dennis et al., 2015) and inevitably go extinct at an exponential rate. Although these models have provided valuable information on extinction probabilities and critical thresholds, they may not adequately capture the nature of environmental variability in many systems. In reality, environmental fluctuations in nature are bounded and may be more often large than small (d’Onofrio, 2013; Fieberg and Ellner, 2001; McLaughlin et al., 2002; Silva et al., 1991; Stacey and Taper, 1992). This fundamental difference raises an important question. What is the effect of bounded environmental fluctuations on populations experiencing a strong Allee effect? To answer this question, I analyze ordinary differential equation (ODE) models of populations experiencing Allee effects and bounded fluctuations in their demographic rates. Unlike previous studies that often focus exclusively on demographic stochasticity (Allen et al., 2005; Ackleh et al., 2007) or use diffusion approximations, this work models environmental variation using a continuous-time Markov chain with any number of states representing different environmental conditions experienced by the population. The environmental state determines the demographic parameters of the ODEs. Hence, one gets population dynamics driven by a Markovian environment, i.e., a piecewise deterministic Markov process (PDMP) (Davis, 1984). PDMPs allow for modeling environmental regimes with potentially large but bounded fluctuations, making them increasingly valuable for ecological applications (Benaïm and Lobry, 2016; Hening and Strickler, 2019; Hening and Nguyen, 2020; Hening et al., 2021; Benaïm et al., 2023; Monmarché et al., 2024).

My analysis reveals several key findings. First, even when populations can persist in each of the environmental states individually, a demographic mismatch between the environmental states can result in inevitable extinction. Second, when population persistence is possible, there emerge two distinct Allee thresholds rather than a single critical density. For populations whose density lies below the lower threshold, extinction is inevitable. When the population density lies above the higher threshold, the population persists indefinitely. Most interestingly, for populations between the two thresholds, extinction or persistence occur with complementary, non-zero probabilities–creating a zone of probabilistic outcomes not captured by deterministic models.

The analysis further explores how the speed at which the environment switches between states influences these probabilities of extinction, revealing that the relationship between environmental fluctuation rates and extinction risk is more complex than previously understood. To demonstrate ecological applications, these analytical findings are applied to specific scenarios of biological importance: populations experiencing mate limitation and predator satiation.

## Models and Methods

### The stochastic model of Allee dynamics

Consider a population with density *N* living in a randomly fluctuating environment. The traditional approach to modeling environmental stochasticity in continuous-time is to formulate the model using stochastic differential equations (Foley, 1994; Lande et al., 2003; Dennis, 2002). However, environmental stochasticity in these models is given by a Brownian motion on log densities. This form of stochasticity has no auto-correlation structure, and fluctuations can be arbitrarily large over arbitrarily small time intervals.

In contrast, here environmental fluctuations *E*(*t*) are modeled as a continuous-time Markov chain with any number (*k*) of states. The environment transitions from state *i* to state *j* at a rate *λ*_*ij*_, that is, ℙ [*E*(*t* + Δ*t*) = *j*|*E*(*t*) = *i*] ≈ *λ*_*ij*_Δ*t* for small Δ*t >* 0. This approach to modeling the environment ensures that the fluctuations are bounded, allows for auto-correlations, and provides great flexibility in the long-term distribution of environmental effects. I assume that this Markov chain is irreducible and, consequently, there is a unique stationary distribution *π* = (*π*_1_, … , *π*_*k*_), where *π*_*i*_ represents the long-term probability of the environment being in state *i*.

The per-capita growth rate *f*_*E*_(*N*) of the population is completely dependent on the environmental state *E* and the population density *N* . The population dynamics are given by a piece-wise deterministic Markov process (PDMP), which was introduced by Davis (1984) and has been used extensively to model stochastic dynamics in a variety of disciplines:

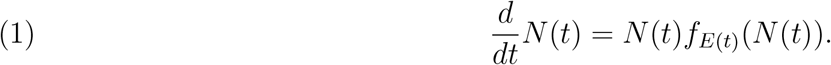

In each environmental state, the population exhibits an Allee effect. Specifically, for the environmental state *E* = *i*, there is an Allee threshold *A*_*i*_ and a carrying capacity *K*_*i*_ such that *f*_*i*_(*A*_*i*_) = *f*_*i*_(*K*_*i*_) = 0, *f*_*i*_(*N*) *<* 0 for *N < A*_*i*_ or *N > K*_*i*_, and *f*_*i*_(*N*) *>* 0 for *A*_*i*_ *< N < K*_*i*_. Thus, in this environmental state, the population rapidly declines to extinction whenever its initial density is below *A*_*i*_. In contrast, the population asymptotically approaches its carrying capacity *K*_*i*_ when its initial density is above *A*_*i*_.

The irreducibility of the Markov chain determining the environmental dynamics allows us to define the mean-field model given by:

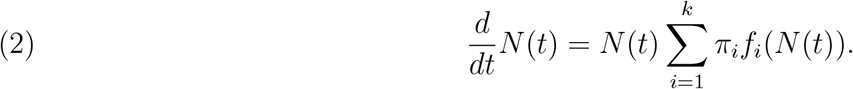

### Analytical and Numerical Methods

In Appendix Appendix, the dynamics of (1), which describes the population growth in a fluctuating environment, are classified by identifying the initial conditions that lead to long-term extinction or persistence and the probabilities of these outcomes. Extinction for an initial condition (*E*(0), *N* (0)) = (*i, n*) means 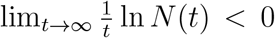 with non-zero probability. In other words, with non-zero probability, the population density decreases exponentially over time, approaching zero. Alternatively, persistence means there exists a minimal density *m >* 0 such that *N* (*t*) ≥ *m* for sufficiently large *t* and with non-zero probability. The mathematical arguments mainly rely on classical results about ordinary differential equations and probability.

Numerical simulations are implemented using a hybrid of Gillespie (1977)’s method for exact simulations of Markov chains and an ordinary differential equation (ODE) solver. Specifically, Gillespie’s method is used to determine the times *T*_1_ *< T*_2_ *<* … at which the environmental state *E*(*t*) changes and the environmental states *E*(*T*_1_), *E*(*T*_2_), … at these times. During each interval [*T*_*i*_, *T*_*i*+1_), the ODE (1) with *E* = *E*(*T*_*i*_) and the initial condition 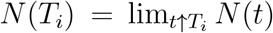 is solved numerically using the deSolve package in R (Soetaert et al., 2010).

## Results

### A Dynamical Dichotomy

The first main finding is a fundamental dichotomy in the dynamical behavior of (1) (Fig. 1, Appendix). This dichotomy depends on three key quantities: the minimum of the Allee thresholds <u>*A*</u> = min_*i*_ *A*_*i*_, the maximum of the Allee thresholds *Ā* = max_*i*_ <u>*A*</u>_*i*_, and the minimum of the carrying capacities <u>*K*</u> = min_*i*_ *K*_*i*_. When the maximal Allee threshold *Ā* is greater than the minimal carrying capacity <u>*K*</u> (a filled circle is to the left of an unfilled circle in Fig. 1A), a population at its carrying capacity in one environment state will eventually find itself below the Allee threshold in another environmental state, and decline. While this decline may be followed by increases in population densities due to favorable environmental shifts, the population always asymptotically approaches extinction, i.e., lim_*t*→∞_ *N* (*t*) = 0 with probability one (Fig. 1B). When *Ā > K*, I say there is a *demographic mismatch* between the environmental states.

**Figure 1.**
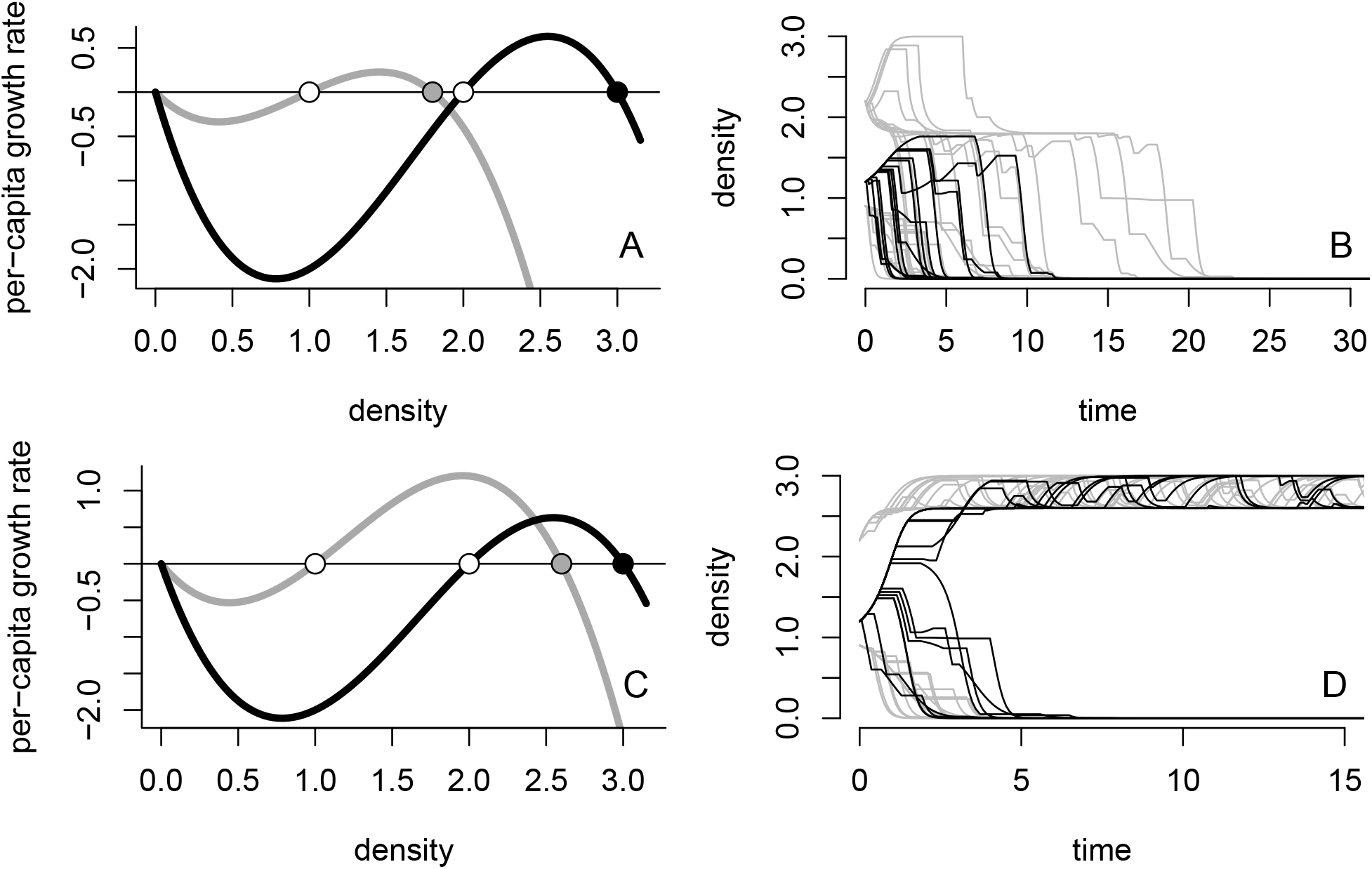
A dynamical dichotomy for populations experiencing strong Allee effects in an environment randomly switching between two states (*k* = 2). In panels A and C, the per-capita growth rates for both environmental states are plotted with Allee thresholds *A*_*i*_ as unfilled circles and carrying capacities *K*_*i*_ as filled circles. In B and D, multiple simulations for three initial densities: one below <u>*A*</u> (light gray), one above *A* (light gray), and one between <u>*A*</u> and *Ā* (black). In A and B, there is demographic mismatch (*<u>K</u> < A*) and the populations always tend toward extinction, i.e., a filled circle lies to the left of an unfilled circle. In C and D, population persistence is possible as *<u>K</u> > A*, i.e., all filled circles lie to the right of the unfilled circles. Furthermore, persistence or extinction occurs with complementary non-zero probabilities for the initial condition lying between <u>*A*</u> and *Ā*. Parameters: *f*_*i*_(*N*) = (*N*− *A*_*i*_)(*K*_*i*_− *N*) with transition rates *λ*_12_ = *λ*_21_ = 1, *A*_1_ = 1, *A*_2_ = 2, and *K*_2_ = 3. For panels A and B, *K*_1_ = 2.6; for panels C and D, *K*_1_ = 1.8.

When there is no demographic mismatch (*Ā < K* as in Fig. 1C), the minimum and maximum Allee thresholds determine the long-term fate of the population (Fig. 1D). If the initial population density is below the minimum Allee threshold (*N* (0) *< <u>A</u>*), then the population continuously decreases to extinction at an exponential rate (lower, gray simulation curves in Fig. 1D). If the initial population density is above the maximum Allee threshold (*N* (0) *> Ā*), then the population density always remains above this threshold and the population persists, that is, *N* (*t*) ≥*Ā* for all time *t* ≥0 (upper, gray simulation curves in Fig. 1D). If the initial population density lies between the minimum and maximum Allee thresholds (*<u>A</u> < N* (0) *< A*), then the long-term fate of the population is stochastic: with non-zero probability, the population goes asymptotically extinct and with a complementary non-zero probability, it persists (black simulation curves in Fig. 1D). This stochastic outcome is determined by the specific sequence of environmental states experienced by the population in the early stages of its dynamics, before its density can either fall below <u>*A*</u> or rise above *Ā*.

### Extinction risk depends on the speed of the environmental dynamics

For initial densities *N* (0) between the minimum and maximum Allee thresholds, the non-zero probabilities of extinction versus persistence depend on how quickly the environment switches between different states. Specifically, suppose that the environmental transition rates are of the form *sλ*_*ij*_ where *s >* 0 determines the speed at which the transitions occur. Although the speed *s* does not affect the fraction *π*_*i*_ of time spent in each environmental state *i*, they have a strong effect on the probabilities of extinction versus persistence (Fig. 2, Proposition 1 in Appendix). Specifically, when transition rates are slow (ie *s* ≈0 but positive), the probability of extinction strongly depends on the initial population density *N* (0) and the initial environmental state *E*(0) = *i*. If *N* (0) *< A*_*i*_, then the probability of extinction is close to 1, otherwise the probability of extinction is close to 0. Intuitively, if the environmental states are changing slowly, then the Allee threshold of the initial environmental state determines, with high likelihood, the long-term population fate (compare the lighter curves in Fig. 2A versus B). Alternatively, if environmental dynamics is fast (*s* ≫0), then the Allee threshold *A*_*_ of the mean field dynamics of equation (2) determines the fate of the population (darker curves in Fig. 2). If the population density is initially below *A*_*_, the probability of extinction is high; otherwise, it is low. Although the Allee threshold for the mean field *A*_*_ always lies between the minimum and maximum Allee thresholds, it is generally not the environmental average ∑_*i*_ *π*_*i*_*A*_*i*_ of the Allee thresholds for the different environments. This stems from Jensen’s inequality, which implies that for a non-linear function, the average of the function values may differ from the function of the average. In this case, the non-linear relationship between environmental conditions and Allee thresholds causes this discrepancy.

**Figure 2.**
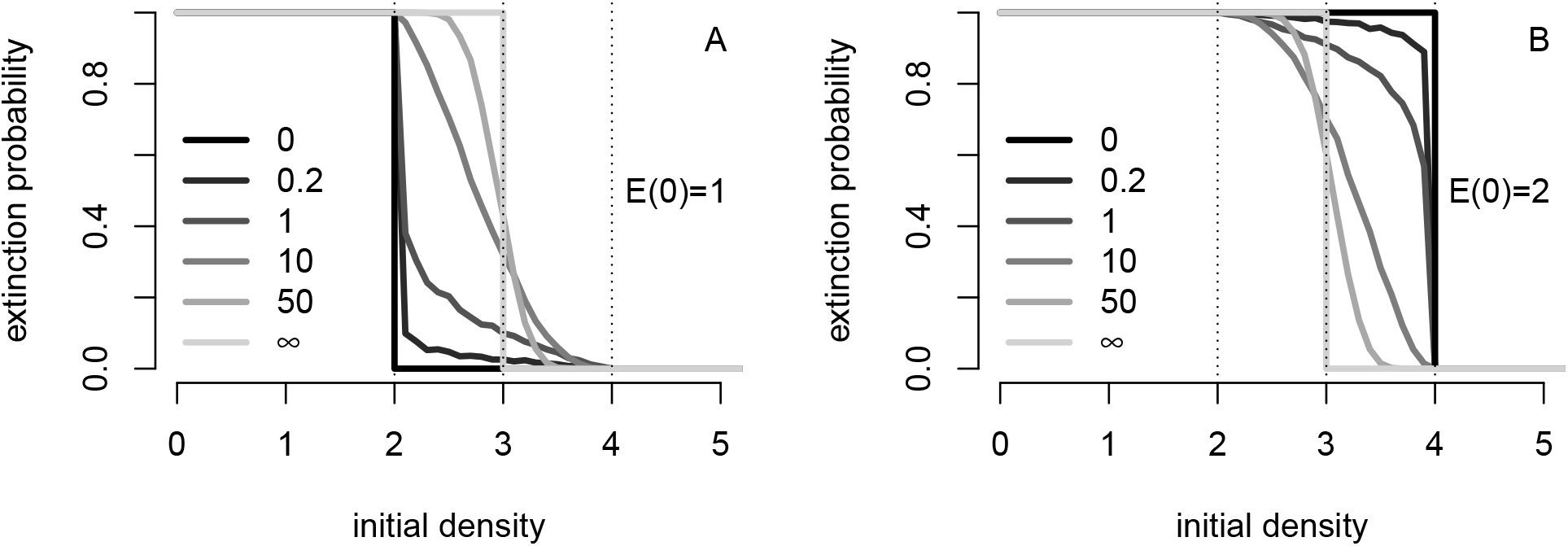
Environmental switching rates alter extinction probabilities. The probability of going asymptotically extinct as a function of the initial population density *N* (0) and different initial environmental states *E*(0). Different shaded curves correspond to different transition rates *λ*_12_ = *λ*_21_. Dashed lines from left to right correspond to minimal Allee threshold *A*, the Allee threshold *A*_*_ of the mean model, and the maximal Allee threshold *Ā*. Parameters: *f*_*i*_(*N*) = (*N*− *A*_*i*_)(*K*_*i*_− *N*) with transition rates *λ*_12_ = *λ*_21_ as shown, *A*_1_ = 2, *A*_2_ = 4, and *K*_1_ = *K*_2_ = 5.

### Simple environmental variation never leads to demographic mismatch

We say that environmental variation is simple if the environmental effects have a one-dimensional parameterization, say *e*, and this parameter has a monotonic effect on the per-capita growth rate. More specifically, assume that all per-capita growth rates are of the form *f*_*i*_(*N*) = *f* (*N, e*_*i*_) where either *f* is non-decreasing in *e* for all *N* , or *f* is non-increasing in *e* for all *N* . For example, increasing *e* always increases birth rates and lowers mortality rates, or increasing *e* always decreases birth rates and increases mortality rates. The environmental variation of this form (Lemma 1 in Appendix) never creates a demographic mismatch. Hence, provided there exist positive Allee thresholds in all environments, simple environmental variation cannot lead to unconditional extinction i.e. all initial conditions lead to asymptotic extinction with probaiblity one.

### Demographic mismatch via complex environmental variation

Complex environmental variation, however, can lead to demographic mismatch. To illustrate how, we examine specific models that separately account for mate limitation and predator satiation. In these models, like most demographic models, there are parameter values for which the per-capita growth is negative for all densities and, consequently, would lead to extinction. We call environments with these parameter values inhospitable environments. For these situations, extinction always occurs whenever one of the environments is inhospitable even if other environments have non-zero Allee thresholds *A*_*i*_ and carrying capacities *K*_*i*_ (Proposition 2 in Appendix).

To model mate limitation, let *N/*(*h*+*N*) be the probability that an individual is mated where *h* is the population density resulting in 50% of individuals being mated (Dennis, 1989; Boukal and Berec, 2002).

Mated individuals produce offspring at a rate *b*. All individuals experience density-independent mortality *d* and density-dependent mortality *cN* where *c* measures the strength of intraspecific competition. In environment *i*, the per-capita growth rate equals

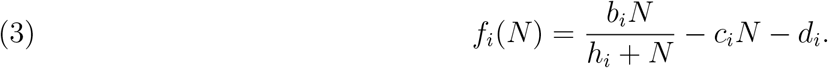

This per-capita growth rate has a positive Allee threshold *A*_*i*_ and a positive carrying capacity *K*_*i*_ whenever 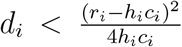 where *r*_*i*_ = *b*_*i*_ − *d*_*i*_ is the intrinsic rate of growth in the absence of mate limitation. Whenever this inequality is not satisfied (i.e. density-independent mortality is too high), environments are inhospitable.

The environmental variation in any one of the demographic parameters, *b, h, d*, or *c*, in equation (3) corresponds to simple environmental variation and consequently does not cause a demographic mismatch. However, variation in multiple parameters can lead to demographic mismatch when some environments have low carrying capacities and other environments have large Allee thresholds. Specifically, this creates a situation where the minimum carrying capacity <u>*K*</u> falls below the maximum Allee threshold *Ā*. For example, when one environment has weaker density-dependent mortality (i.e., smaller *c*_*i*_) and greater mate limitation (i.e., larger *h*_*i*_) than another environment, then demographic mismatch can occur (Fig. 3A).

**Figure 3.**
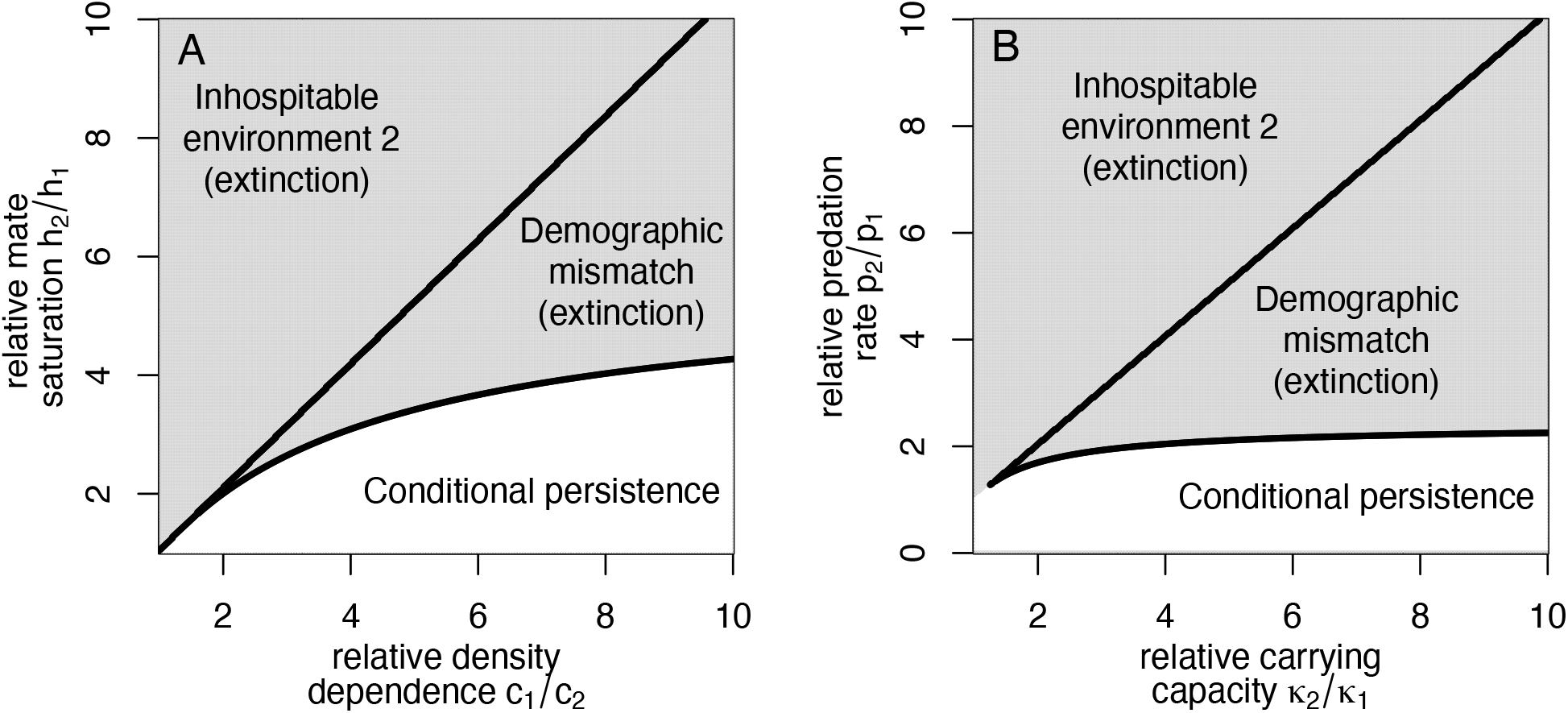
Demographic consequences of mate-limitation (A) or predator satiation (B) in a fluctuating environment. Regions of parameter space correspond to three types of ecological dynamics: conditional persistence in which there is persistence or extinction depending on initial conditions; demographic mismatch in which the maximal Allee threshold *Ā* is greater than the minimal carrying capacity <u>*K*</u> and the population is ultimately driven extinct; and an inhospitable environmental state in which the per-capita growth rate is negative for all densities and the population ultimately goes extinct.

To model a prey population experiencing predation from a generalist predator, we use the model of Noy-Meir (1975). In the absence of predation, the population exhibits Logistic dynamics with intrinsic rate of growth *r* and carrying capacity *κ*. The maximal per-capita predation rate is *p* with a half saturation prey density *h*. The per-capita growth rate in the environmental state *i* is

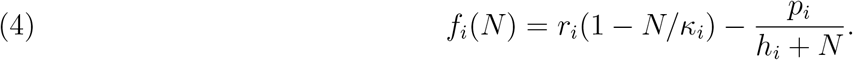

In environment *i*, there is an Allee threshold *A*_*i*_ and carrying capacity *K*_*i*_ if 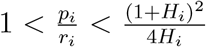 where *H*_*i*_ = *h*_*i*_*/κ*_*i*_. The first inequality ensures a negative per-capita growth rate at low population densities, and the second inequality ensures the existence of *A*_*i*_ and *K*_*i*_. When the second inequality is violated 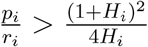-the maximal predation rate is too high), the environment is inhospitable.

If the environmental states only affect any one of the demographic rates (i.e. *r, κ, h* or *p*), then the environmental variation is simple and there is no demographic mismatch. However, simultaneous variation in the maximal predation rate *p* and the prey’s carrying capacity *κ* can lead to demographic mismatch. When the prey carrying capacity *κ* is small, the carrying capacity *K* in the presence of predation is low. Alternatively, if the prey carrying capacity *κ* is high and the maximal predation *p* is high (but not too high), then there is a high Allee threshold *A*. Environments that alternate between these conditions—specifically when one environment produces a low *K* value and another environment produces a high *A* value such that *<u>K</u> < Ā* —create a demographic mismatch that leads to inevitable extinction (Fig. 3B).

## Discussion

In nature, environmental fluctuations are bounded and may be typically large rather than small. However, these realities are not accommodated by stochastic differential equations that are commonly used to model environmental fluctuations in populations with overlapping generations (May, 1973; Dennis, 1989; Foley, 1994; Lande et al., 2003; Nolting and Abbott, 2016; Hening and Nguyen, 2018). Here, environmental stochasticity is captured by Markovian dynamics between any number of states. These Markovian dynamics are inherently bounded and allow for arbitrarily complex environmental dynamics including a predominance of large rather than small fluctuations. I have shown that, for populations experiencing a strong Allee effect, this additional environmental realism can dramatically impact the long-term population dynamics. Unlike stochastic differential equation models that predict strong Allee effects always drive population extinct (Dennis et al., 2015), models allowing for bounded, stochastic fluctuations generate two distinct scenarios depending on the relationship between environmental states.

The first scenario occurs when there is a demographic mismatch between environments: namely, when the carrying capacity in one environment lies below the Allee threshold in another environment. Even when environmental conditions individually allow for conditional persistence of the population, fluctuating between these conditions may lead to unavoidable extinction. Demographic mismatch only occurs when environmental shifts have non-monotonic impacts on the per-capita growth rate—where shifts increase growth rates over some density ranges but decrease them over others. For example, in populations experiencing mate limitation, mismatch can occur when environmental conditions that reduce intraspecific competition simultaneously increase mate limitation. Temperature fluctuations may generate such trade-offs. The metabolic theory of ecology predicts that intraspecific competition increases at higher temperatures (effectively lowering carrying capacity) (Savage et al., 2004)—a prediction supported by experimental evidence (Bernhardt et al., 2018). In contrast, a recent metaanalysis suggests that higher temperatures may enhance mating success (Pilakouta and Baillet, 2022), potentially lowering Allee thresholds. When these effects occur together, temperature fluctuations can create the conditions for demographic mismatch. Similarly, the analysis presented here suggests that environments that support higher prey carrying capacities while simultaneously enabling greater maximal predation rates can create a demographic mismatch for prey populations with predator-driven Allee effects. This positive relationship between prey carrying capacities and predation intensity aligns with general scaling laws showing that total predator biomass increases with total prey biomass in diverse terrestrial and aquatic communities (Hatton et al., 2015; Perkins et al., 2022). Kaul et al. (2016) showed that Allee thresholds for the bacterivore *Cafeteria roenbergensis* depend in a complex manner on the availability of carbon resources and predation levels, potentially illustrating these types of demographic trade-offs.

The second scenario reveals a novel finding: in the absence of demographic mismatch, populations experiencing environmental fluctuations and Allee effects can persist, an important difference from classical SDE models that predict inevitable extinction (Dennis, 2002; Dennis et al., 2015). In these cases, there are two critical Allee threshold densities *<u>A</u> < Ā* rather than just one. If the initial population density lies below the lower threshold density <u>*A*</u>, the population always goes extinct. If the initial density exceeds the higher threshold density *Ā*, the population always persists and remains above this higher threshold. Similar to deterministic models (Dennis, 1989; Boukal and Berec, 2002; Schreiber, 2003), the fate of populations with initial densities outside the interval [*<u>A</u>, Ā*] is deterministic. However, unlike deterministic models, populations with initial densities within this interval have stochastic fates: they either go extinct or persist with complementary, non-zero probabilities.

In the absence of demographic mismatch, the rate at which the environment fluctuates between different states determines the sensitivity of the results to the initial conditions. The dynamics of a system with fast environmental switching can be closely approximated by the mean-field model (2). This deterministic model has a single critical threshold density *A*^*^ that lies between the two threshold densities of the stochastic model (*<u>A</u> < A*^*^ *< Ā*). Hence, with fast switching, extinction becomes highly likely (though not certain) when the initial density falls between <u>*A*</u> and *A*^*^, while persistence becomes highly likely when the initial density lies between *A*^*^ and *Ā*. In contrast, when environmental switching is slow, the outcomes depend heavily on both the initial population density and the initial environmental state. Specifically, if the initial population density falls below the Allee threshold of the initial environmental state, extinction becomes highly likely but not certain. Conversely, when the initial density exceeds the initial Allee threshold, persistence becomes highly likely.

These findings complement and extend previous work on stochastic Allee effects. Liebhold and Bascompte (2003) studied a discrete-time model that incorporated an Allee effect and environmental stochasticity, but unlike our model, theirs lacked negative density-dependence. Their model, parameterized for the North American Gypsy moth pest (*Lymantria dispar*), revealed threshold behavior similar to that observed here. Specifically, when initial densities fell below the critical threshold of their mean-field model, environmental stochasticity increased persistence probability by occasionally pushing populations above the critical density. In contrast, when initial densities exceeded this threshold, stochasticity increased the risk of extinction. Roth and Schreiber (2014) later provided mathematical confirmation of these simulation results. However, important distinctions exist between these studies and the work presented here: these earlier models employed unbounded, log-normal noise similar to SDE approaches, and the probabilistic outcomes they observed stemmed primarily from the absence of negative density-dependence, rather than from the bounded nature of environmental fluctuations.

Our findings suggest several promising directions for future research that incorporate additional biological realism. First, spatial structure in deterministic models with Allee effects generates multiple alternative states corresponding to different configurations of patches with low and high occupancy (Vortkamp et al., 2020); extending the analyses presented here to spatially explicit stochastic models could provide insight into how bounded environmental fluctuations affect these spatial patterns. Second, interspecific interactions could be incorporated, particularly since competition can produce complex frequency-dependent dynamics through mechanisms such as shared social information (Gil et al., 2019) or reproductive interference (Schreiber et al., 2019). Third, eco-evolutionary feedbacks may cause Allee thresholds to shift as population traits evolve (Klausmeier et al., 2020), raising questions about how bounded environmental fluctuations might interact with these dynamic thresholds. Most crucially, our demonstration that bounded environmental stochasticity can enable long-term persistence for populations with Allee effects suggests that conservation strategies for threatened populations may need to reconsider traditional extinction risk assessments that rely on unbounded stochastic models. Collectively, this work highlights the importance of incorporating realistic environmental variation when predicting the fate of populations experiencing Allee effects, a critical consideration for both theoretical ecology and conservation biology.

## Appendix. Mathematical Proofs of Main Results

In this Appendix, we provide precise statements and mathematical proofs of results concerning (1). Let ℝ = ( −∞,∞) denote the real line and ℝ_+_ = [0, ∞) the non-negative reals. For our main result, we need the following assumptions:

**A1:** The per-capita growth rate functions *f*_*i*_ : ℝ_+_ → ℝ are Lipschitz.

**A2:** For each *i*, there exists *K*_*i*_ *> A*_*i*_ *>* 0 such that *f*_*i*_(*A*_*i*_) = *f*_*i*_(*K*_*i*_) = 0, *f*_*i*_(*x*) *>* 0 for *x*∈ (*A*_*i*_, *K*_*i*_), and *f*_*i*_(*x*) *<* 0 for *x*∈ [0, *A*_*i*_) ∪ (*K*_*i*_, *∞*).

**A3:** The finite state Markov chain *E*(*t*) is irreducible with stationary distribution *π* = (*π*_1_, *π*_2_, … , *π*_*k*_)

### Theorem 1.

*Assume* ***A1****–****A3***. *Define A* = min_*i*_ *A*_*i*_, *Ā* = max_*i*_ *A*_*i*_, <u>*K*</u> = min_*i*_ *K*.

*If Ā > <u>K</u>, then* lim_*t*→∞_ *N* (*t*) = 0 *with probability one whenever N* (0) *>* 0.

*If Ā < <u>K</u> and Ā* ≠ *<u>A</u>, then*

i. *N* (*t*) ≥ *Ā for all t* ≥ 0 *whenever N* (0) ≥ *Ā*,
ii. *a*(*x, i*) := ℙ [*N* (*t*) ≥ *Ā for some t* ≥ 1|(*N* (0), *E*(0)) = (*x, i*)] *>* 0 *and* ℙ [lim_*t*→∞_ *N* (*t*) = 0|(*N* (0), *E*(0)) = (*x, i*)] = 1 − *a*(*x, i*) *>* 0 *whenever x* ∈ [*A, Ā*) *and* 1 ≤ *i* ≤ *k, and*
iii. lim_*t*→∞_ *N* (*t*) = 0 *whenever N* (0) ∈ (0, *A*).

Note: On the event {lim_*t*→∞_ *N* (*t*) = 0}, one can show that 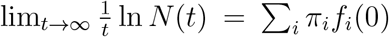 with probability one.

*Proof*. Let *ϕ*_*i*_(*t, x*) to denote the flow of 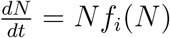 in environment *i* i.e. *t* ↦ *ϕ*_*i*_(*t, x*) is the solution Of 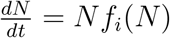 satisfying *N* (0) = *x*.

The proof makes repeated use of Lévy 0-1 Law. To state this Law in our context, for any integer *m* ≥ 0, define ℱ_*m*_ to be the *σ*-algebra generated by the random variables

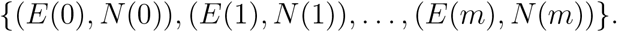

ℱ_0_, ℱ_1_, … is in an increasing sequence of *σ*-algebras. Define the *σ*-algebra ℱ_∞_ as ∩_*m*≥1_ℱ_*m*_. Given any set 𝒜 ∈ ℱ_∞_, let 𝟙_𝒜_ : ℱ_*m*_ →{0, 1} be the indicator function, i.e. equals 1 if the event 𝒜 has occurred and 0 otherwise.

**Theorem** (Lévy 0-1 Law). *Let* 𝒜 ∈ ℱ_∞_. *Then* lim_*m*→∞_ 𝔼 [𝟙_𝒜_ |ℱ_*m*_] = 𝟙_A_.

Now, assume that *Ā > <u>K</u>*. As *f*_*i*_(*x*) *<* 0 for all *x* ≥ *K, N* (*t*) eventually enters and remains in the interval 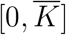. Hence, without loss of generality, assume that *N* (0) lies in this interval. As *f*_*i*_(*x*) *<* 0 for all *x < <u>A</u>*, lim_*t*→∞_ *N* (*t*) = 0 whenever *N* (*t*) enters the interval [0, <u>*A*</u>). Define 𝒜 = {*N* (*k*) *< <u>A</u>* for some *k* ≥ 1} which lies in ℱ_∞_.

Given 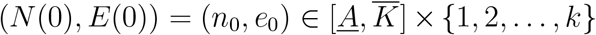, I will show that there is a positive probability that *N* (*t*) enters [0, *A*). To see why, define *i*^*^ = arg max_*i*_ *A*_*i*_ and *i*_*_ = arg min_*i*_ *K*_*i*_. Choose *t*^*^ *>* 0 such that (i) 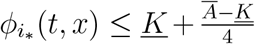 for all 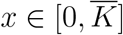 and *t* ≥ *t*^*^, and (ii) 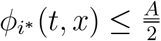 for all 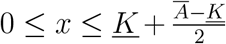 and *t* ≥ *t*^*^. This choice of *t*^*^ is possible as (i) *f*_*i*_ (*x*) *<* 0 for 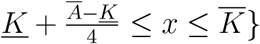 and (ii) 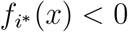 for 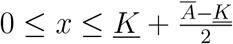 . Choose *t*_*_ *>* 0 such that 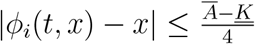 for all *i*, 0 ≤ *t* ≤ *t*_*_, and 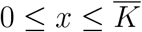.

This choice of *t*_*_ is possible as *ϕ*_*i*_ is continuous and 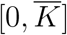 is compact.

For any *m* ≥ 0, let *T*_1_(*m*) = inf{*t* ≥ *m* : *E*(*t*) = *i*_*_} be the first time after *t* = *m* that the environmental state is *i*_*_. Let *T*_2_(*m*) = inf{*t* ≥ *T*_1_(*m*) : *E*(*t*) ≠*i*_*_} be the first time the environmental state leaves the state *i*_*_ after *T*_1_(*m*). On the event {*T*_2_(*m*) − *T*_1_(*m*) *> t*^*^}, the definition of *t*^*^ implies that *N* (*T*_2_(*m*)) ≤ <u>*K*</u> + (*Ā* − *K*)*/*4. Define *T*_3_(*m*) = inf{*t* ≥ *T*_2_(*m*) : *E*(*t*) = *i*^*^} to be the first time the environmental state is *i*^*^ after *T*_2_(*m*). On the event {*T*_2_(*m*) − *T*_1_(*m*) *> t*^*^} ∩ {*T*_3_(*m*) − *T*_2_(*m*) *< t*_*_}, the definitions of *t*_*_, *t*^*^ imply that *N* (*T*_3_(*m*)) ≤ *K* + (*Ā* − *K*)*/*2 *< A*. Define *T*_4_(*m*) = inf{*t* ≥ *T*_3_(*m*) : *E*(*t*) ≠ *i*^*^} to be the first time after *t* = *T*_3_(*m*) to leave the state *i*^*^. On the event

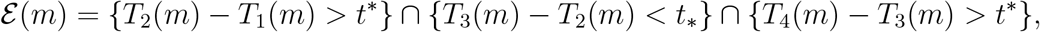

we have *N* (*T*_4_(*m*)) *< A/*2. In particular, ℰ (*m*) ⊂ 𝒜. Let

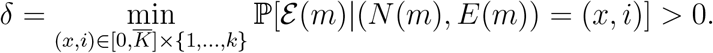

As ℰ (*m*) ⊂ 𝒜, 𝔼 [𝟙 _𝒜_| ℱ_*m*_] ≥ 𝔼 [𝟙 _ℰ (*m*)_| ℱ _*m*_] ≥ *δ*. By the Lévy 0-1 Law, 𝟙 _𝒜_ = 1 with probability one. Namely, lim_*t*→∞_ *N* (*t*) = 0 with probability one.

Now, assume that *Ā < K*. As *f*_*i*_(*x*) *<* 0 for all (*x, i*) ∈ (0, *A*) × {1, 2, … , *k*}, lim_*t*→∞_ *N* (*t*) = 0 whenever *N* (0) *< A*. As *f*_*i*_(*x*) *>* 0 for all (*x, i*) ∈ (*Ā, K*) × {1, 2, … , *k*}, *N* (*t*) ≥ *Ā* whenever *N* (0) *> Ā*.

Assume *N* (0) ∈ (*A, Ā*). (Re)define *Ã* = (*A* + *Ā*)*/*2, *i*_*_ = arg min_*i*_ *A*_*i*_ and *i*^*^ = arg max_*i*_ *A*_*i*_. Choose *t*_1_ *>* 0 and *ε >* 0 such that 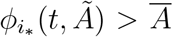 and 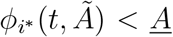 for all *t* ≥ *t*_1_. Define the stopping times *T*_*_(*m*) = inf{*t* ≥ *m* : *E*(*t*) = *i*_*_}, *T*_**_ = inf{*t* ≥ *T*_*_(*m*) : *E*(*t*) ≠ *i*_*_}, *T* ^*^(*m*) = inf{*t* ≥ *m* : *E*(*t*) = *i*^*^}, and *T* ^**^(*m*) = inf *t T* ^*^(*m*) : *E*(*t*) ≠ *i*^*^ }. Define the event 𝒜 = *N* (*t*) *∉* [*A, Ā*] for some *t* ≥ 0}. Choose *δ >* 0 such that

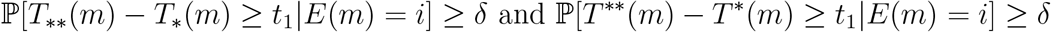

for all environmental states *i* and *m* ≥ 1. It follows that for all *m* ≥ 1

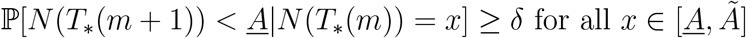

and

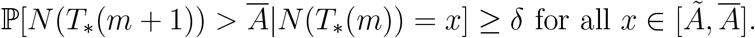

For all *m* ≥ 1,

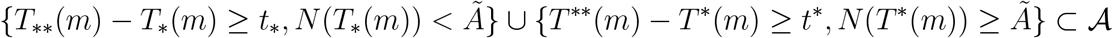

Hence, by the Lévy 0-1 Law, 𝟙 _𝒜_ = 1 almost surely. In particular, for *x* ∈ (*A, Ā*) and 1 ≤ *i* ≤ *k*, ℙ [lim_*t*→∞_ *N* (*t*) = 0|(*N* (0), *E*(0)) = (*x, i*)] = 1 − *a*(*x, i*) where *a*(*x, i*) := ℙ[*N* (*t*) ≥ *A* for some *t* ≥ 1|(*N* (0), *E*(0)) = (*x, i*)].

Finally, I need to show that 0 *< a*(*x, i*) *<* 1 for all *x* ∈ (*A, Ā*). Assume *N* (0) = *x* ∈ (*A, Ā*) and *E*(0) = *i*. Choose *t*_1_ *>* 0 and *ε >* 0 such that *ϕ*_*i*_(*t, x*) ∈ (*A* + *ε, Ā* − *ε*) for *t* ∈ [0, *t*_1_]. Choose *t*_2_ *>* 0 such that 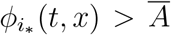; for all *t ≥ t*_2_ and *x* ∈ [*A* + *ε, Ā*− *ε*]. Let *T*_1_ = inf *t* ≥ 0 : *E*(*t*) = *i*_*_ and *T*_2_ = inf {*t* ≥*T*_1_ : *E*(*t*) ≠ *i*_*_} . Then *a*(*x, i*) ≥ℙ [ {*T*_1_ ≤*t*_1_, *T*_2_ ≥ *t*_2_ }] *>* 0. The fact that *a*(*x, i*) *<* 1 is proved in a similar way choosing *t*_2_ such that 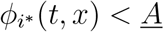.

The next proposition provides limiting values for the probabilities *a*(*x, i*) in Theorem 1 when switching between environments occurs quickly or slowly.

### Proposition 1.

*Assume* ***A1****–****A3*** *hold and Ā < K and Ā* ≠ *A. For the Markov chain* {*E*(*t*)}, *assume the transition rates for i*≠ *j are of the form λ*_*ij*_ = *τα*_*ij*_ *where τ >* 0 *and α*_*ij*_ ≥ 0. *For each τ >* 0, 1 ≤ *i* ≤ *k, and x* ∈ [*A, Ā*], *define a*_*τ*_ (*x, i*) := ℙ [*N* (*t*) ≥ *Ā for some t* ≥ 1|(*N* (0), *E*(0)) = (*x, i*)].*Then*

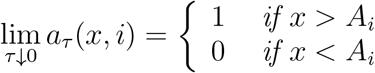

*Furthermore, if there exists a unique A*_*_ ∈ [*A, Ā*] *such that ∑* _*i*_ *π*_*i*_*f*_*i*_(*A*_*_) = 0, *then*

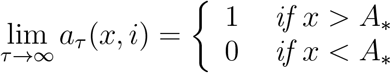

*Proof*. Let {*N*_*τ*_ (*t*), *E*_*τ*_ (*t*)} be the Markov chain with speed *τ >* 0.

To prove the limit in the case of *τ* ↓ 0, let *i, x > A*_*i*_, and *ε >* 0 be given. Choose *t*_*i*_ *>* 0 be such that *ϕ*_*i*_(*t*_*i*_, *x*) ≥ *Ā* + (*K*_*i*_ − *Ā*)*/*2. Assume (*N* (0), *E*(0)) =(*x, i*) and let *T* = inf{*t* ≥ 0 : *E*(*t*) ≠ *i*}. Choose *τ >* 0 sufficiently small so that ℙ [*T < t*_*i*_] = exp(−*τ ∑*_*j*≠ *i*_ *λ*_*ij*_*t*_*i*_) *< ε*. It follows that *a*_*τ*_ (*x, i*) ≥ ℙ [*N* (*t*_*i*_) *> Ā*] ≥1 −*ε*. As *ε >* 0 was arbitrary, lim_*τ*↓0_ *a*_*τ*_ (*i, x*) = 1 as claimed. The proof that lim_*τ*↓0_ *a*_*τ*_ (*i, x*) = 0 for *x < A*_*i*_ follows from a similar argument.

To prove the limit in the case of *τ* ↓ 0,let *ϕ*_*_(*t, x*) be the flow of the averaged vector field

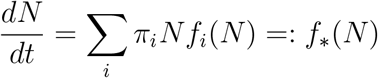

i.e. *t* ⟼ *ϕ*_*_(*t, x*) is the unique solution to this differential equation with initial condition *N* (0) = *x*. Let *A*_*_, by assumption, be the unique equilibrium in (*<u>A</u>, Ā*). By continuity *f*_*_(*N*) *<* 0 for *N < A*_*_ and *f*_*_(*N*) *>* 0 for *N* ∈ (*A*_*_, <u>*K*</u>). Let (*x, i*) ∈ (*A*_*_, *Ā*) × {1, 2, … , *k*} be given. Assume that (*N* (0), *E*(0)) = (*x, i*). Choose *T >* 0 such that 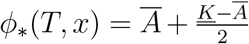 . Benaïm and Strickler (2019a, Lemma A.4) implies that for all *δ >* 0

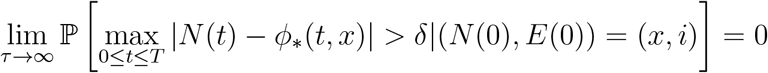

Choosing 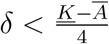 implies lim_*τ*→∞_ *a*_*τ*_ (*x, i*) = 1 as claimed. The proof that lim_*τ*↓0_ *a*_*τ*_ (*i, x*) = 0 for *x < A*_*_ follows from a similar argument.

The next lemma implies that “simple” environments can not result in a demographic mismatch.

### Lemma 1.

*Let f* (*x, y*) *be function such that for all y there exist K*(*y*) *> A*(*y*) *>* 0 *such that f* (*A*(*y*), *y*) = *f* (*K*(*y*), *y*) = 0 *for all y, f* (*x, y*) *>* 0 *for A*(*y*) *< x < K*(*y*), *and f* (*x, y*) *<* 0 *for x* ∈ (0, *A*(*y*)) ∪ (*K*(*y*), ∞). *If y ⟼f* (*x, y*) *is non-decreasing for all x, then A*(*y*) *is a non-increasing function and K*(*y*) *is a non-decreasing function*.

*Proof*. Choose any *y*_2_ *> y*_1_. As *f* (*x, y*_2_) ≥ *f* (*x, y*_1_) *>* 0 for all *x* ∈ (*A*(*y*_1_), *K*(*y*_1_)), *A*(*y*_2_) ≤ *A*(*y*_1_) and *K*(*y*_2_) ≥ *K*(*y*_1_).

Finally, in the cases where there is no Allee effect or there is an inhospitable environment, (Benaïm and Strickler, 2019b, Theorems 3.1 and 3.2) implies lim_*t*→∞_ *N* (*t*) = 0 whenever ∑_*i*_ *π*_*i*_*f*_*i*_(0) *<* 0 and *N* (*t*) is stochastically persistent whenever ∑_*i*_ *π*_*i*_*f*_*i*_(0) *>* 0.

